# Codon usage bias in prokaryotic genomes and environmental adaptation

**DOI:** 10.1101/2020.04.03.023309

**Authors:** Davide Arella, Maddalena Dilucca, Andrea Giansanti

## Abstract

In each genome synonymous codons are used with different frequencies; this phenomenon is known as codon usage bias. The preferred codons tend to correspond to the most highly expressed tRNAs. It had been known that codon usage bias can influence the cellular fitness and that might be associated with the lifestyle of the organism. To test the impact of environments on genome evolution we studied the relationship between codon usage bias and the phenotypic traits of 615 prokaryotic organisms. Principal component analysis revealed that prokaryotes with a specific phenotypic characteristic and living in similar environmental conditions have similar codon preferences, accessed by the Relative Synonymous Codon Usage (RSCU), and a similar tRNA availability gauged by the tRNA gene copy number (tGCN). In addition, by measuring the average tRNA adaptation index (tAI) for each genome, we discovered that organisms able to live in multiple habitats, including facultative organisms, mesophiles and pathogenic bacteria, exhibit lower extents of translational efficiency, consistent with their need to adapt to different environments.

This is the first large-scale study that examines the role of translational efficiency in the environmental adaptation of prokaryotes. Our results show that synonymous codon choices might be under strong translational selection, adapting the codons to the tRNA pool to different extents depending on the organism’s lifestyle needs.

## INTRODUCTION

The genetic code is degenerate, i.e., some amino acids are encoded by more than one codon. Although coding for the same amino acid, synonymous codons are not equally used, a phenomenon known as codon usage bias (or shorter, codon bias) [14]. Codon usage might differ widely not only between organisms, but also within a genome and within a single gene [17,28]. A lot of factors might cause different codon usage bias and the selective forces influencing it, such as selection for optimized translation, expression and location in genes, rate of evolution, secondary structure, nucleotide composition, protein length and environment [33]. It was demonstrated that many bacteria and yeast undergo translational selection, with highly expressed genes preferentially using codons assumed to be translated faster and/or more accurately by the ribosome [13, 3]. Thus, the codon usage bias within a genome usually reflects the selection pressure for translational optimization of highly expressed genes. The choice of preferred codons in a single genome is most closely correlated with abundance of the cognate tRNA molecules [3,18,19,9] and further influenced by the genome’s GC content [7, 16].

It was introduced by Ardersson and Kurland [1] and then substantiated by Kudla *et al*. [23] that selection towards highly adapted codons in highly expressed proteins has a global effect on the cell, resulting in an increase in cellular fitness. This suggest that the global extent of codon usage bias of an organism might be associated with its phenotypic traits. Following this idea, Botzman *et al*. determines an association between the lifestyles of several prokaryotic organisms and variations in their codon usage [5]. Their results indicated that species living in a wide range of habitats have low codon usage bias, which is consistent with the need to adapt to different environments. Furthermore, by analyzing 11 diverse microbial community sequencing samples, Roller *et al*. demonstrats that microbes living in the same ecological niche share a common preference for codon usage, regardless of their phylogenetic diversity [32]. Complementing these studies, the analysis of acidophilic bacteria reveales that they preferentially have low codon bias, consistent with both their capacity to live in a wide range of habitats and their slow growth rate [15].

The physical requirements that are optimal for prokaryotic growth vary dramatically for different types of bacteria and archea. As a group, bacteria display the widest variation of all organisms in their ability to inhabit different environments. One of the most prominent differences between prokaryotes is their requirement for, and response to, atmospheric oxygen (O_2_). On the basis of oxygen requirements, bacteria can be divided into obligate aerobes (they have absolute requirement for oxygen in order to grow), obligate anaerobes (they grow only in the absence of oxygen), facultative anaerobes (they thrive in the presence of oxygen but can grow in its absence), aerotolerant anaerobes (they do not use oxygen but are indifferent to the presence of oxygen) and microaerophiles (they require a minimum level of oxygen for growth, about 1%–10%). Prokaryotes have adapted to a wide range of temperatures.

The National Center for Biotechnology Information (NCBI) Microbial Genome Project Database uses five terms to categorize the temperature range an organism grows at, where cryophilic refers to -30° to -2° C, psychrophilic refers to -1° to 10° C, mesophilic refers to 11° to 45° C, thermophilic refers to 46° to 75° C, hyperthermophilic refers to above 75° C, and organisms that live at ranges that overlap with more than one category are labeled as the one corresponding to the largest overlap [42]. Water is a fundamental requirement for life. Some organisms prefer salty environments and are thus called halophiles. There is a wide range of halophilic microorganisms belonging to the domains Bacteria and Archaea. Halophiles are categorized as slight, moderate, or extreme, by the extent of their salinity preference. Slight halophiles prefer 0.3 to 0.8 M (1.7 to 4.8% seawater is 0.6 M or 3.5%), moderate halophiles 0.8 to 3.4 M (4.8 to 20%), and extreme halophiles 3.4 to 5.1 M (20 to 30%) salt content [26]. In relation to habitat conditions, some bacteria are able to thrive in a wide variety of environmental conditions, while others can thrive only in a narrow range of environmental conditions.

The aim of our study is to calculate the codon usage bias of more than 600 prokaryotic species and to better understand the role of codon usage bias in the adaptation of prokaryotes to the environment.

## MATERIALS AND METHODS

We analyzed the extent of codon usage bias in 615 organisms (544 bacteria and 71 archaea) reported in Supplementary Material (see Table S1 of Excel file). Classification of environment and characteristics were downloaded from the Additional file 2 of the work conducted by Botzman *et al*. [5]. Nucleotide sequences were downloaded from the FTP server of the National Center for Biotechnology Information [4]. The tRNA gene copy number for each organism was retrieved from the Genomic tRNA database (GtRNAdb) [24] (available at the site http://gtrnadb.ucsc.edu). The number of organisms annotated with each property is detailed in Table 1.

**Table 1.**
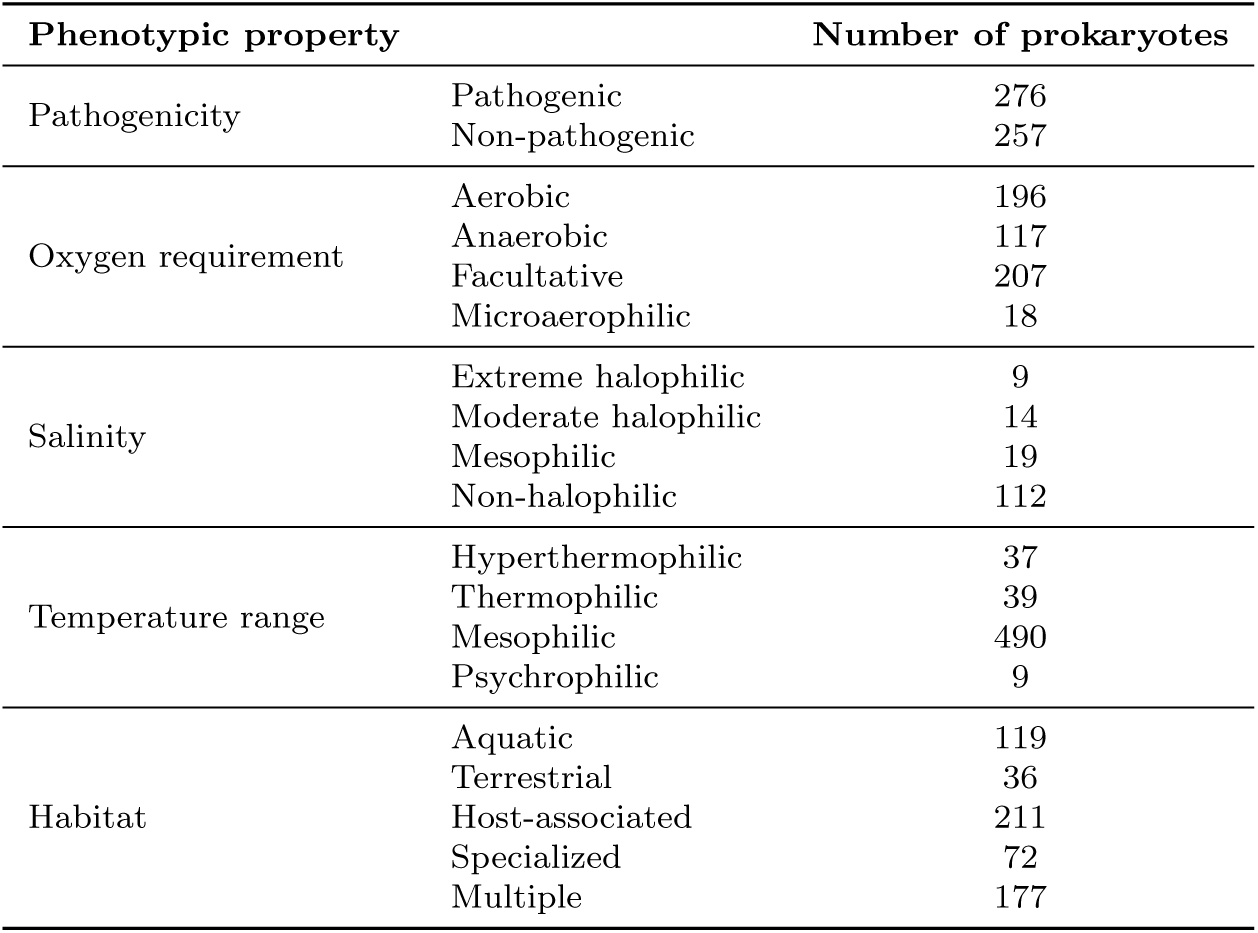
Classification of the prokaryotes in the dataset by phenotypic traits.

To detect different patterns of codon usage among the genes of a species, heat-maps were drawn with the CIMminer software [41] (http://discover.nci.nih.gov/cimminer).

### RSCU calculation

The Relative Synonymous Codon Usage (RSCU) [35] is the observed frequency of a codon divided by the expected frequency if all the synonymous codons were used equally. The RSCU is computed for each codon of each amino acid and it is formally defined as follows. Let *n*_*i*_ denote the number of synonymous codons encoding for the amino acid *i* (codon degeneracy) and let *X*_*ij*_ denote the number of occurrences of the codon *j* for amino acid *i*. The RSCU for codon *j* encoding the amino acid *i* is defined as

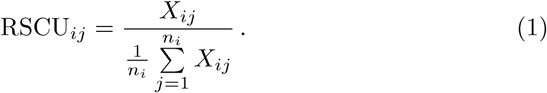

RSCU is a real value comprised between 0 and the number of synonymous codons for that amino acid, i.e., *n*_*i*_. For average synonymous codon usage (no codon bias) the RSCU is 1. For codon usage more infrequent than the average codon usage, the RSCU is less than one, and for more frequent usage than the average for the amino acid, the RSCU is greater than 1.

We calculated RSCU with a Python script homemade. The RSCU values of the various codons can be considered as the 61 components (excluding the stop codons TAA, TAG and TGA – which are differently used by different species) of vectors which measure codon usage bias in a given gene.

For each genome we calculated the average vector of RSCU, 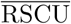 and the similarity between the RSCU vector of a gene and the average RSCU vector of the genome. As a measure of similarity we used the cosine similarity with the following formula:

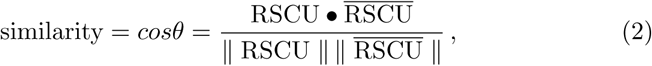

where •denotes the scalar product and ‖RSCU‖ is the norma of the RSCU vector. When the cosine similarity is 1 the two vectors have the same orientation, whereas if the cosine similarity is 0 they are orthogonal to each other.

### PCA

Principal component analysis (PCA) [21] is a multivariate statistical method to transform a set of observations of possibly correlated variables into a set of linearly uncorrelated variables (called principal components) spanning a space of lower dimensionality. The transformation is defined so that the first principal component accounts for the largest possible variance of the data, and each succeeding component in turn has the highest variance possible under the constraint that it is orthogonal to (i.e., uncorrelated with) the preceding components.

We used this technique on the space of 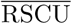 values, where each organism of the dataset was represented as a 61-dimensional vector with the codons as coordinates. The eigenvectors of the associated correlation matrix, ordered according to the magnitude of the corresponding eigenvalues, are the principal components of the original data.

The PCA was performed using the open source software gretl (http://gretl.sourceforge.net). We projected in the plane of the first two principal components all genomes of the dataset. Centroids were calculated as mean value with relative error bars as standard deviation of the mean.

We then carried out an other PCA using the number of tRNA gene copies (tGCN) provided by the GtRNAdb, in order to consider the availability of tRNA for each prokaryote.

### tAI calculation

The speed of protein synthesis is bound to the waiting time for the correct tRNA to enter the ribosomal A site [39], and thus depends on tRNA concentrations [36]. The consequent adaptation of codon usage to tRNA availability [18, 19] is at the basis of tRNA adaptation index (tAI) [30, 10]. It takes advantage of the fact that the tRNA gene copy number across many genomes has a high and positive correlation with tRNA abundance within the cell [18, 27, 22, 11]. The tAI follows the same mathematical model of CAI [34] – defining for each codon *i* its absolute adaptiveness (*W*_*i*_):

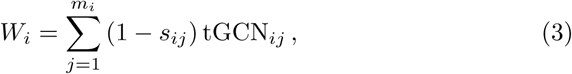

where *m*_*i*_ is the number of tRNA isoacceptors that recognize the *i*th codon, tGCN_*ij*_ is the gene copy number of the *j*-th tRNA that recognizes the *i*-th codon and *s*_*ij*_ is a selective constraint on the efficiency of the codon-anticodon coupling. From the *W*_*i*_ values the relative adaptiveness value *w*_*i*_ of a codon is obtained as

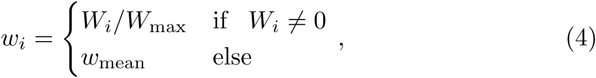

where *W*_max_ is the maximum *W*_*i*_ value and *w*_mean_ is the geometric mean of all *w*_*i*_ with *W*_*i*_ ≠ 0. Finally, the tRNA adaptation index tAI_*g*_ of a gene *g* is computed as the geometric mean of the relative adaptiveness values of its codons

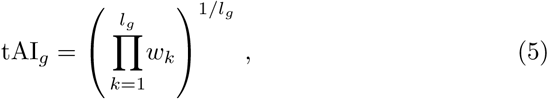

where *k* is the codon defined by the *k*-th triplet in gene *g* and *l*_*g*_ is the length of the gene in codons (except the stop codon). The critical issue for tAI is the selection of a meaningful set of *s*_*ij*_ values, i.e., weights that represent the efficiency of the interactions between codons and tRNAs. Assuming that tRNA usage is maximal for highly expressed genes, these values are chosen in order to optimize the correlation of tAI values with expression levels.

We calculated tAI values using the tAI package provided by Mario dos Reis on GitHub (https://github.com/mariodosreis/tai). This is an R package that implements the tAI as described in dos Reis *et al*. [30].

We divided the prokaryotes into groups according to their environmental characteristics and pathogenicity and then we compared the distributions of the average tAI values belonging to these groups. Mann-Whitney U-test and Kruskal-Wallis H-test were used to verify if the differences between the distributions were statistically significant with *p*-value *<* 0.05. Significance tests were implemented in Python using the statistical functions of the SciPy library (https://www.scipy.org/).

## RESULTS

### RSCU values

Previous observations (see [14, 3, 28]) pointed to the fact that each bacterial species has a specific pattern and level of codon usage bias, which is strongly shared by the majority of its genes; codon bias in specialized categories of genes appears to be just a modulation of the distinctive codon bias of the species [8]. To check this statement, we computed the RSCU values for all the coding regions of each prokaryote in the dataset. In Figure 1 is shown an example of heat-map of RSCU values for each gene (in the example, of *Escherichia coli* K12 substrain MG1655) that shows the existence of a fingerprint of codon bias for this organism. In order to quantify the degree of similarity between the various RSCU vectors and the average RSCU vector, we calculated the cosine similarity between the RSCU vector of each CDS and the 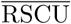 vector belonging to the species (see Figure 2). The similarity distribution showed a mean value near to 1 indicating that for the majority of the CDSs the RSCU vector is similar to the average RSCU vector. Therefore, the 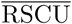 vector is a good descriptor for the codon usage of the whole genome.

**Fig. 1.**
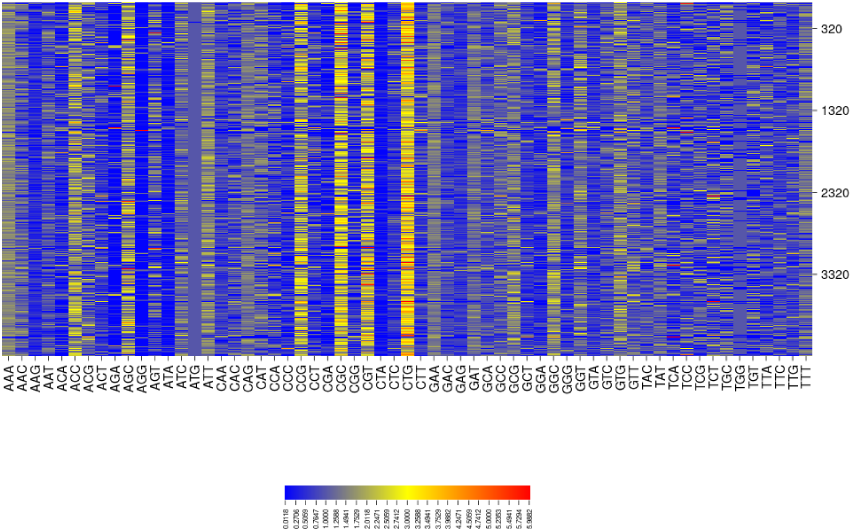
Heat map of RSCU values for each gene of *Escherichia coli* strain K12 substrain MG1655. The 4319 CDSs are in rows and the 61 codons are in columns (in alphabetic order). We note that RSCU vectors of different genes are very similar to each other. So an average RSCU vector can be considered a fingerprint for a species.

**Fig. 2.**
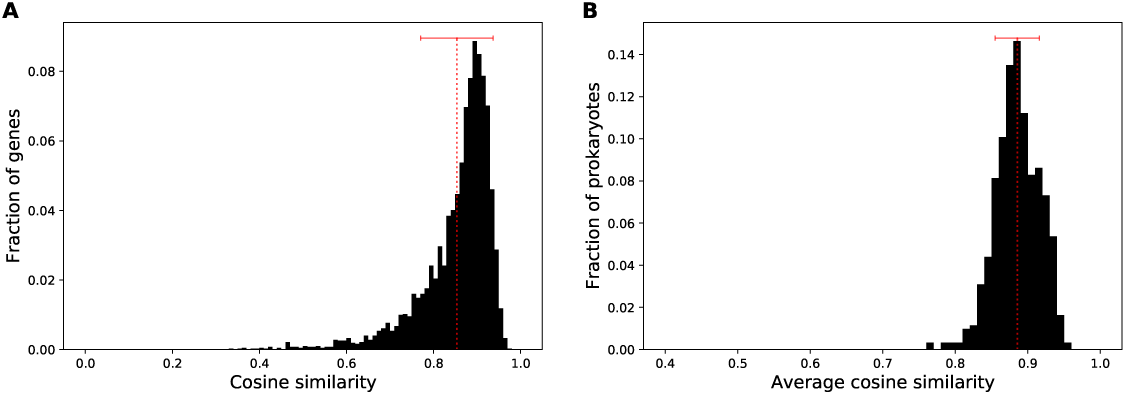
(**A**) Distribution of cosine similarity between 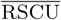 vector and RSCU vectors for all the CDSs of *Escherichia coli* strain K12 substrain MG1655. Mean (0.85) and standard deviation (0.08) of the distribution are plotted in red. (**B**) Relative frequency of the average cosine similarity between RSCU vector of a gene and 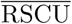 vector of a genome for all the prokaryotes in dataset. Red dotted line denotes the mean (0.88) of the distribution and the error bar the standard deviation (0.03). We note that the average similarity is near to 1, demonstrating that the average RSCU vector is a good descriptor for the codon usage bias of the whole genome.

Overall, this exploration suggests that there should be a strong correlation between codon bias patterns of each species and his evolutionary history. In our opinion, there is an ecological determinant behind this rough classification based on basic codon bias.

### PCA

We analyzed the patterns of synonymous codon usage among the prokaryotes in our dataset using the principal component analysis on the 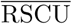 vectors measured for each species (see Figure 3). The two first principal components (PC_1_ and PC_2_) turned out to represent as much as 71% of the total variance of codon bias over the genomes. Interestingly, the distributions of prokaryotes related to different phenotypic characteristics had well separated centroids in this reduced space (four panels of Figure 3). When we characterized the prokaryotes depending on their habitat, the distributions of the organisms exhibited distinct centroids for every habitat (see Figure 4). What we have found represents an important result: prokaryotes with a specific phenotypic characteristic and living in similar environmental conditions demonstrated a similar codon usage bias measured in terms of 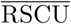 vectors. In other words, if a set of genomes are physically and functionally connected in an environment, their corresponding genes share common codon bias features.

**Fig. 3.**
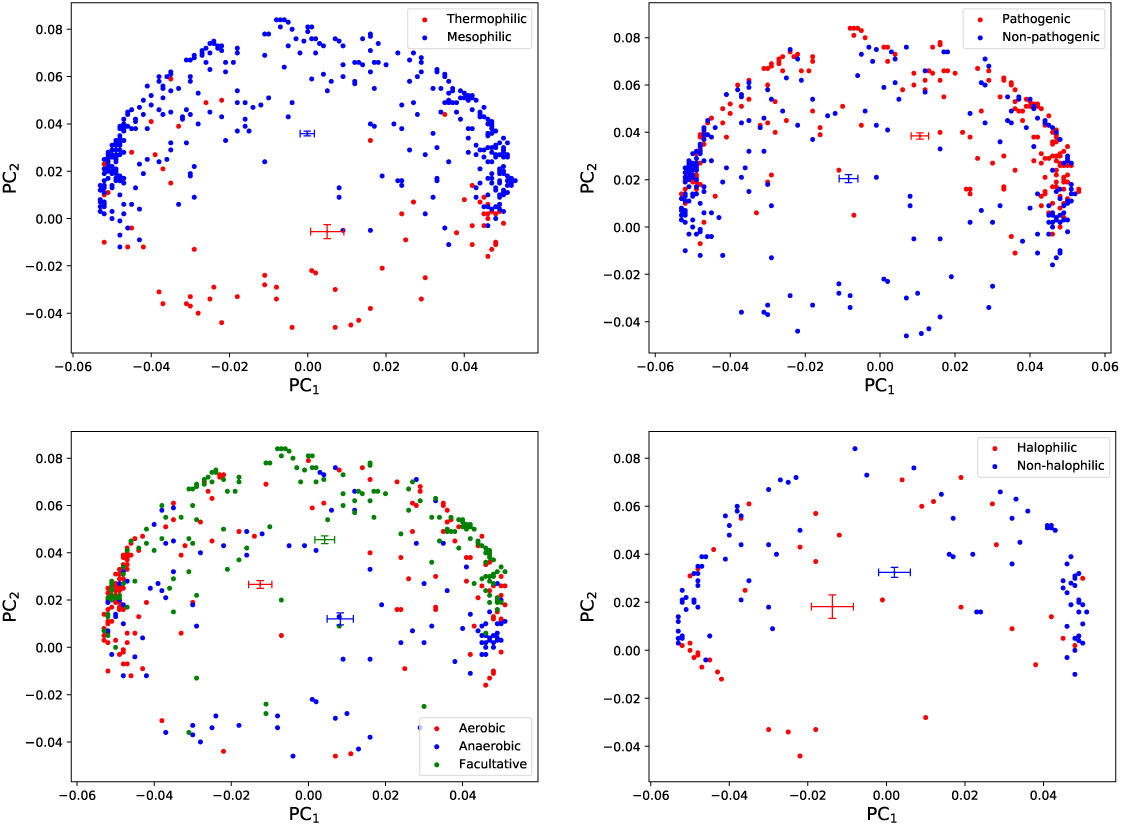
Principal component analysis using 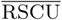 vectors of all the prokaryotes in the dataset. The first principal component (PC_1_) accounted for 51.6% of the total variation and the second principal component (PC_2_) accounted for 19.7% of the total variation. We projected in the PC_1_-PC_2_ plane the organisms according to their phenotypic traits. Centroids were calculated as mean value with relative error bars as standard deviation of the mean. Note that prokaryotes have distributions with very well separated centroids.

**Fig. 4.**
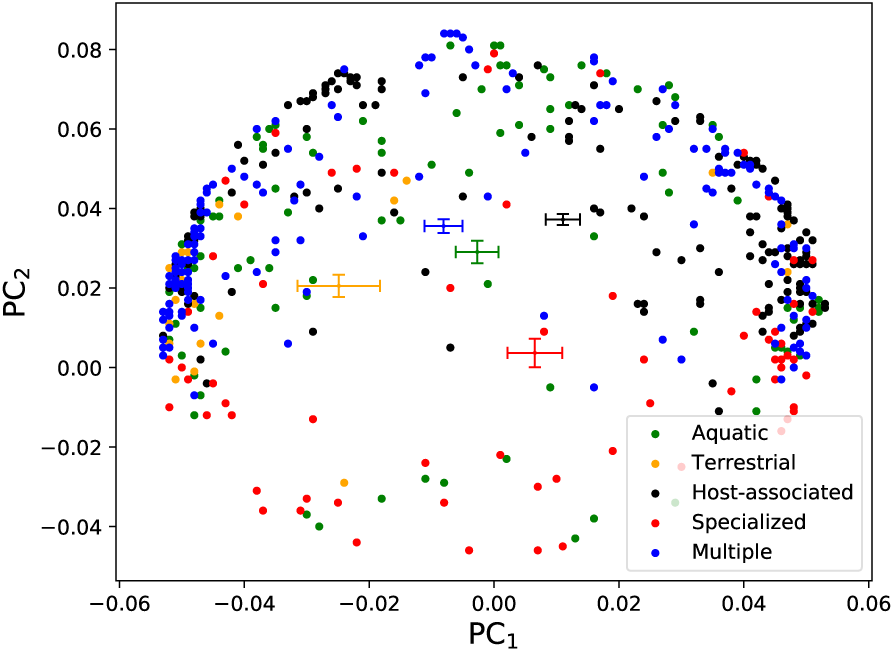
Principal component analysis using 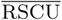 vectors of all the prokaryotes in the dataset. We projected in the PC_1_-PC_2_ plane the organisms according to the habitat where they live. Note that the centroids are well separated.

We carried out another principal component analysis to consider the tRNA availability of the prokaryotic genomes. In order to do that, we used the number of copies of tRNA genes (tGCN) measured for each species in the dataset instead of the 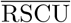 values. Unfortunately, the fraction of the total variance accounted by the first two principal components of this second principal component analysis was only 37%. Therefore, we chose to add the third principal component for the visualization of our dataset, explaining in this way 43% of the total variance.

We divided the prokaryotes into groups according to their lifestyles and plotted them in the space of the first three principal components. Figure 5 shows the distribution of species on the first three principal components of the PCA. As we can see, the centroids of the distributions are well separated in the reduced space of the first three principal components. When we characterized prokaryotes for different habitats (see Figure 6), the distributions of the prokaryotes exhibited distinct centroids. Therefore, the principal component analysis showed us that prokaryotes living in similar physical and chemical constraints share some common feature not only in the codon usage bias but also in the tRNA availability.

**Fig. 5.**
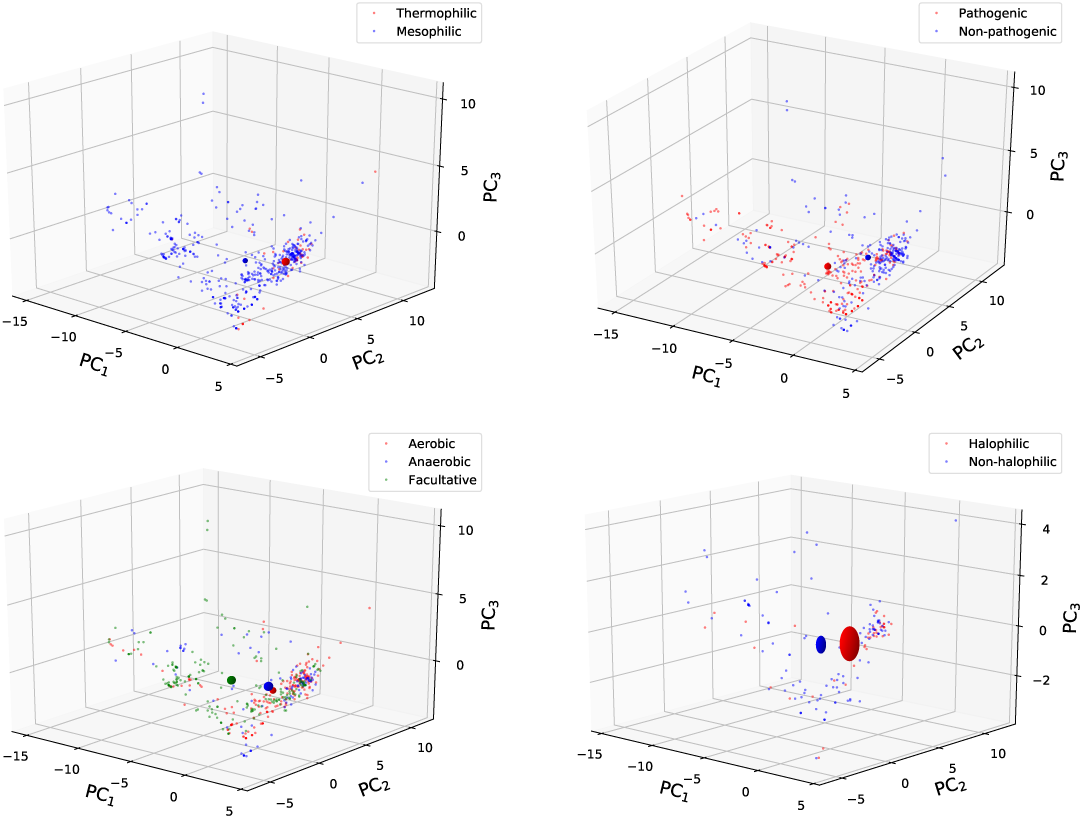
Principal component analysis using tGCN values of all the prokaryotes in the dataset. The first principal component (PC_1_) accounted for 24.7% of the total variation, the second principal component (PC_2_) accounted for 12.6% of the total variation and the third principal component (PC_3_) accounted for 5.5% of the total variation. We projected in the PC_1_-PC_2_-PC_3_ space the organisms according to their phenotypic traits. Centroids of the distributions are represented as spheres centered around mean value and with radius equal the maximum standard deviation of the mean along the axes. Note that prokaryotes have distributions with very well separated centroids.

**Fig. 6.**
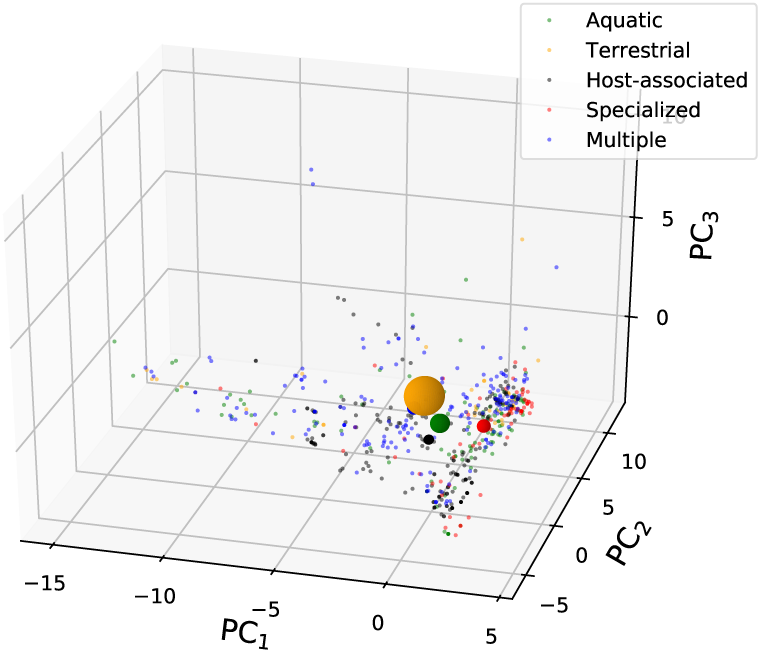
Principal component analysis using tGCN values of all the prokaryotes in the dataset. We projected in the PC_1_-PC_2_-PC_3_ space the organisms according to the habitat where they live. Centroids of the distributions are represented as spheres centered around mean value and with radius equal the maximum standard deviation of the mean along the axes. Note that the centroids are well separated.

### tAI values

As shown in Figure 7, there is a wide distribution of tAI average values across the genomes, ranging between 0.15 and 0.79, with a mean value of 0.39, median of 0.36 and standard deviation of 0.12.

**Fig. 7.**
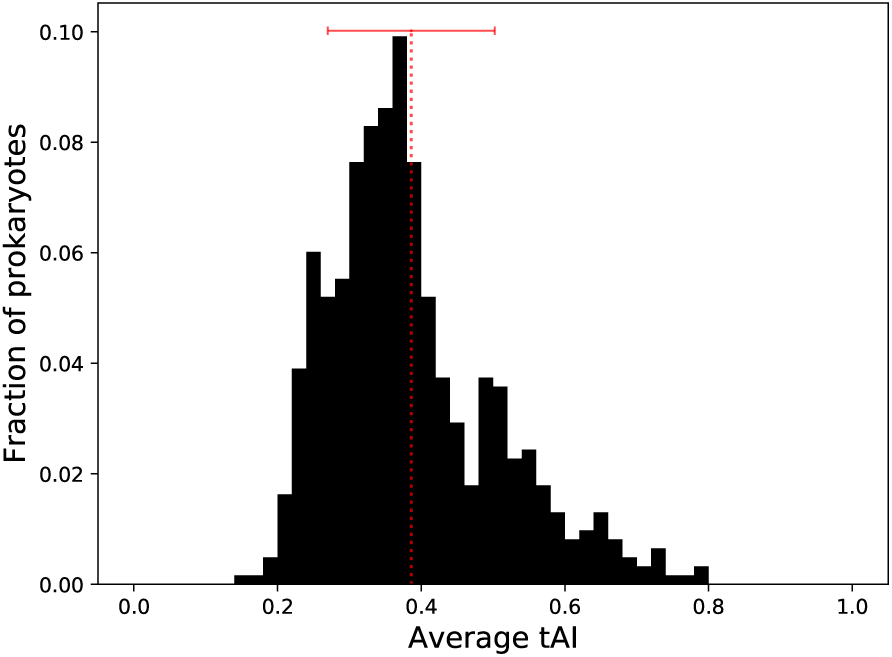
Distribution of 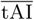 values among the 615 prokaryotes in dataset. Red dotted line denotes mean of the distribution and the error bar the standard deviation.

We annotated each genome with its 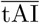 and compared the distributions of the 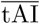 values of different groups of organisms, tagged according to a particular phenotypic trait.

We examined the distribution of 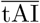 values among thermophilic versus mesophilic prokaryotes (see Figure 8). Organisms that live in different temperature ranges showed statistically significant differences in their 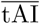 values (*p* = 1.62 × 10^−8^, two-sided Mann-Whitney test): thermophiles demonstrated statistically significantly higher 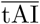 values than mesophiles. As shown in Figure 8, the distribution among pathogenic bacteria is biased to the left compared to non-pathogenic bacteria, with pathogenic prokaryotes having statistically significant lower 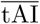 values (*p* = 8.67 × 10^−8^ by two-sided Mann-Whitney test). Groups of prokaryotes classified by their oxygen requirement (Figure 8) differed statistically significantly in the distributions of 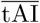 values (*p* = 7.09 × 10^−8^ by Kruskal-Wallis test). Interestingly, facultative organisms exhibited the lowest extent of translational efficiency and their distribution was statistically different from the distributions belonging to the other two groups (*p* = 4.53 × 10^−8^, two-sided Mann-Whitney test between facultative and aerobic; *p* = 1.35 × 10^−4^ between facultative and anaerobic). The difference between aerobic and anaerobic species was not statistically significant (*p* = 0.301 by two-sided Mann-Whitney test). Groups of prokaryotes that live in environments that differ in their salinity levels (Figure 8) did not demonstrate statistically significant differences among them (*p* = 0.161 by two-sided Mann-Whitney test).

**Fig. 8.**
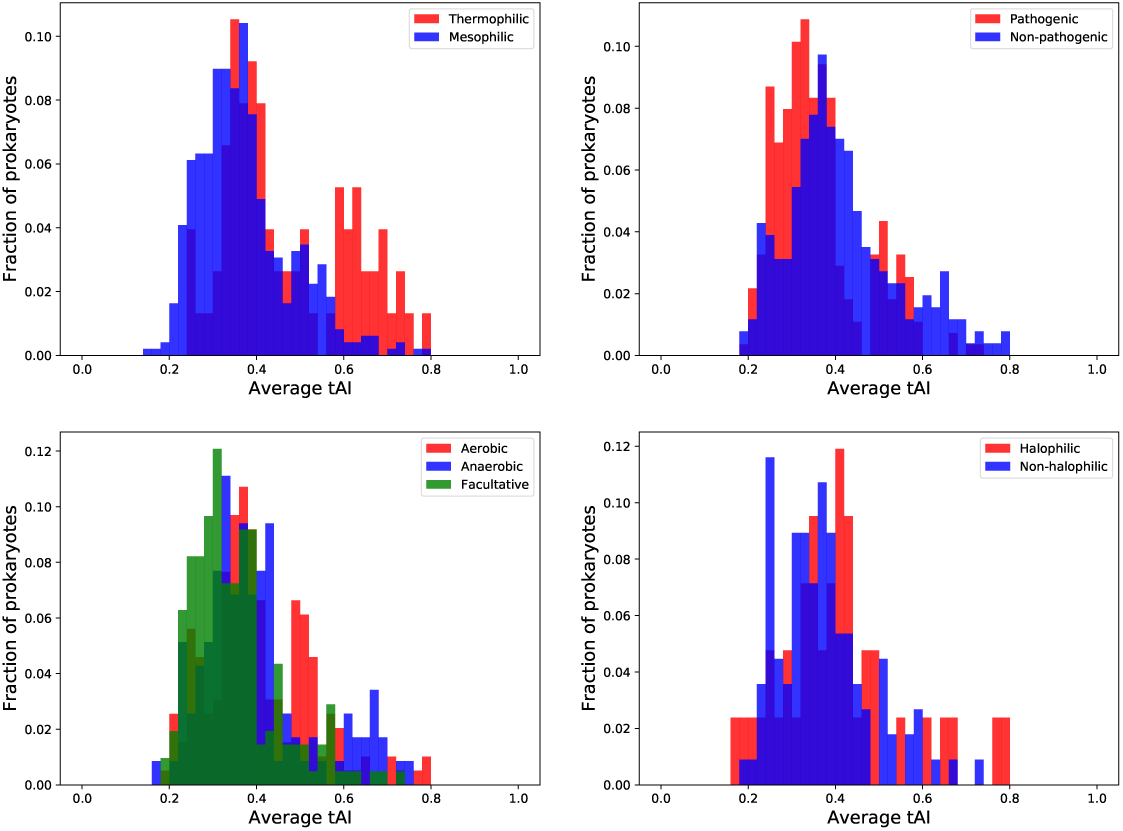
Relative frequency distribution of 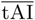 values of organisms classified according to their characteristics.

We then turned to analyze the differences between organisms living in different habitat conditions (see Figure 9). Intriguingly, we found that organisms living in multiple habitats have statistically significant lower 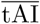 values than organisms living in specialized habitats (*p* = 2.66 × 10^−6^, two-sided Mann-Whitney test). This result is consistent with the results presented above for the other phenotypic traits and generalizes them. Pathogenic bacteria often live in multiple environments outside and within their host, and facultative organisms live in environments with and without oxygen. On the other hand, thermophiles (found above to have a higher extent of translational efficiency) are usually restricted to a specific environment with a specific temperature.

**Fig. 9.**
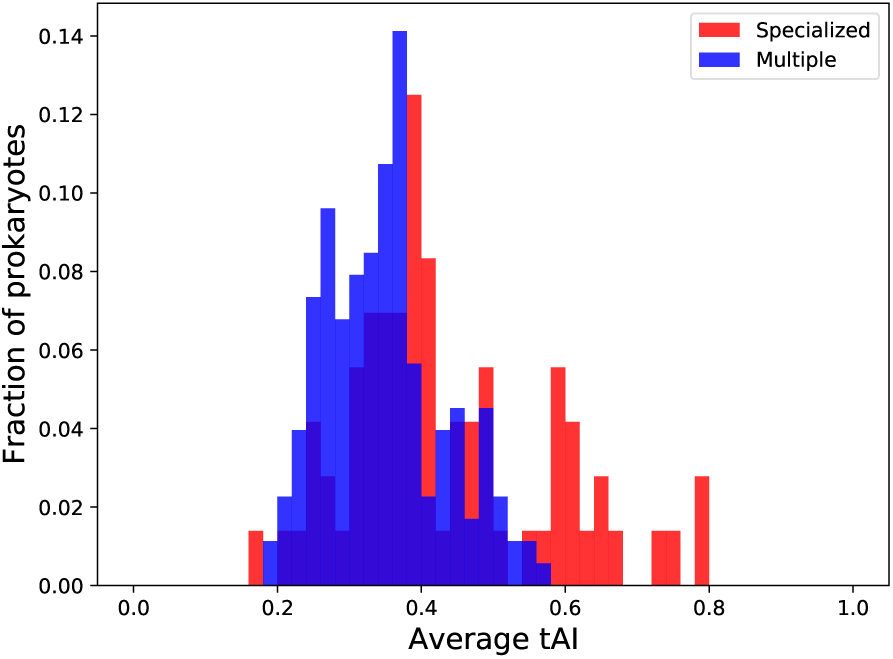
Relative frequency distribution of 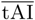 values of prokaryotes divided into two groups: organisms that live in a specialized environment and organisms that live in multiple habitats.

## DISCUSSION

The analysis of codon usage bias has been widely used to characterize both specific and general properties of genes from communities of microorganisms [29]. In this context, we studied the codon usage bias of 615 prokaryotic genomes with optimal codons identified by RSCU values. Each genome was identified by a 61-dimensional 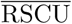 vector that represent, as a fingerprint, the dominating codon bias of the organism. Principal component analysis was applied to 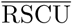 space and confirmed that different phenotypic traits belonging to the species can be identified by codon preferences. These results indicate that similarity in the extents of codon usage bias of different organisms reflects a similar ecological strategy they share.

Previous studies demonstrated the correspondence between ecology preferences and codon adaptation [5, 20]. To stress this idea, we added the information about the adaptation of codon usage to the genomic tRNA gene pool [30, 34] where translational selection is known to be present. It has been observed in several species that *in vivo* concentration of a tRNA bearing a certain anticodon correlates with the number of gene copies coding for this tRNA (for example, in *S. cerevisiae*, Pearsons *r* = 0.91 [27]). This facilitates the investigation of the tRNA pools of any fully sequenced species.

Using the tRNA gene copy numbers, we carried out a PCA of the tRNA repertoire belonging to the 615 prokaryotes. This analysis was less convincing than the previous one (the first three principal components explained only 43% of total variance), but roughly confirmed that different prokaryotic lifestyles can be traced in similarities in the tGCN values.

To enable a comparison of translational efficiency of genes among the species, we calculated the tRNA adaptation index (tAI), an index based on the availability of each of the tRNAs. Several lines of evidence indicate that the tAI-based translation efficiency values are biologically significant and very often higher tAI corresponds to higher protein abundance [30, 25, 38]. Indeed protein expression levels can be artificially increased by designed mutations that increase their codon-tRNA adaptation [27, 38], pointing to a causal relationship between codon usage and expression level.

An organism-scale measure was obtained by computing 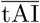, the average of the tAI values of all CDSs in a genome. Our analysis revealed a large variability in this measure: there are organisms showing very high degrees of translation efficiency and organisms exhibiting very low 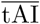 values. The findings of Botzman and Margalit [5] motivated us to compare the distributions of 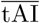 values of groups of prokaryotes with different phenotypic characteristics. Remarkably, we found that the extent of translational efficiency corresponds to the lifestyle of the organism. Even more interestingly, we found that these differences can be explained by considering whether the organisms live in multiple or specialized habitats. We have sustained that organisms that live in a specialized habitat have higher extents of translational adaptation, consistent with their need to adapt efficiently to a specific environment. On the contrary, prokaryotes that live in multiple environments have shown lower 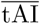 values as they need to be more flexible.

Reasons supporting an evolutionary convergence of codon bias for organisms sharing similar physiology and living in similar habitats, might include the need for a successful exchange of genes by lateral transfer, the sharing of physical parameters such as temperature (preferred amino acids and codons related to thermal adaptation), the sharing of chemical parameters such as nutrient supply (that would differentially affect pyrimidine and purine production), and the sharing of biological parameters such as, for pathogens, the management of genetic variability through codon usage (to escape the host-immune system for instance) [6].

Examples supporting these reasons are several. The bacterium *Aquifex aeolicus*, for instance, occupies the hyperthermophilic niche otherwise dominated by Archaea. After genome analysis, it seems likely that the archaeal genes in *Aquifex* have been introduced by horizontal gene transfer, on top of a typical bacterial gene repertoire, and have been retained owning to the specific selective advantage they provided by enabling the bacterium to thrive in high-temperature habitat [2]. A similar gene transfer has been observed for another hyperthermophilic bacterium, *Thermotoga maritima* [6]. Furthermore, communities of microbes have been shown to share similar tRNA pools to facilitate horizontal gene transfer [37], which also implies a limited choice of preferred codons that are cognate to the shared community tRNA pool. This is consistent with the findings of the present work.

Freilich *et al*. showed that most bacterial organisms choose one of two alternative ecological strategies: living in multiple habitats with a large extent of co-habitation, associated with a typically fast rate of growth or living in a specialized niche with little co-habitation, associated with a typically slow rate of growth [12]. Independently, Rocha demonstrated that fast growing bacteria have more tRNA genes of fewer types and suggested that the translation in those organisms depends on fast tRNA diffusion to the ribosome [31, 40]. Our findings tie these two results together and suggest that organisms may adjust to metabolic variability and competition by maintaining a low extent of adaptation of their genes to the tRNA pool (reflected by their low 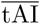 values).

## CONCLUSIONS

It is generally accepted that the speed at which ribosomes decode a codon depends on the cellular concentration of the tRNA that recognize it. In many genomes, it has been shown that the preferred codons are those that tend to match the most abundant tRNAs, suggesting that selection for optimal translation is a main factor influencing codon usage bias.

It has been recognized that codon usage bias can affect the cellular fitness and that might be associated with the lifestyle of the organism. To test this hypothesis we studied the relationship between codon usage bias and the phenotypic traits of 615 prokaryotic organisms. Principal component analysis revealed that prokaryotes with a specific phenotypic characteristic and living in similar environmental conditions have similar codon preferences, accessed by the Relative Synonymous Codon Usage (RSCU), and a similar tRNA availability gauged by the tRNA gene copy number (tGCN). Furthermore, by measuring the average tAI for each genome, we discovered that organisms able to live in a wide range of habitats exhibit lower extents of translational efficiency, consistent with their need to adapt to different environments. Pathogenic prokaryotes also demonstrate lower degree of translational efficiency than non-pathogenic prokaryotes, in accord with the multiple environments that many pathogens occupy. Facultative organisms, which are able to growth in the presence or in the absence of oxygen, have lower values of average tAI. Lower extent of translational efficiency was observed also in mesophiles which are capable of growing in a wide range of temperatures.

We point out that this is the first large-scale study that examines the role of translational efficiency in the adaptation of prokaryotes to the environment in which they live. Our results show that synonymous codon choices might be under strong translational selection, adapting the codons to the tRNA pool to different extents depending on the organism’s lifestyle needs.

## References

1. Andersson, S., Kurland, C.: Codon preferences in free-living microorganisms. Microbiology and Molecular Biology Reviews 54(2), 198–210 (1990)

2. Aravind, L., Tatusov, R.L., Wolf, Y.I., Walker, D.R., Koonin, E.V.: Evidence for massive gene exchange between archaeal and bacterial hyperthermophiles. Trends in Genetics 14(11), 442–444 (1998)

3. Bennetzen, J.L., Hall, B.D.: Codon selection in yeast. Journal of Biological Chemistry 257(6), 3026–3031 (1982)

4. Benson, D.A., Cavanaugh, M., Clark, K., Karsch-Mizrachi, I., Lipman, D.J., Ostell, J., Sayers, E.W.: GenBank. Nucleic acids research 41 (D1), D36–D42 (2012)

5. Botzman, M., Margalit, H.: Variation in global codon usage bias among prokaryotic organisms is associated with their lifestyles. Genome biology 12(10), R109 (2011)

6. Carbone, A., Kepes, F., Zinovyev, A.: Codon bias signatures, organization of microorganisms in codon space, and lifestyle. Molecular biology and evolution 22(3), 547–561 (2005)

7. Chen, S.L., Lee, W., Hottes, A.K., Shapiro, L., McAdams, H.H.: Codon usage between genomes is constrained by genome-wide mutational processes. Proceedings of the National Academy of Sciences 101(10), 3480–3485 (2004)

8. Dilucca, M., Cimini, G., Semmoloni, A., Deiana, A., Giansanti, A.: Codon bias patterns of E. colis interacting proteins. PloS one 10(11) (2015)

9. Dong, H., Nilsson, L., Kurland, C.G.: Co-variation of tRNA abundance and codon usage in Escherichia coli at different growth rates. Journal of molecular biology 260(5), 649–663 (1996)

10. Dos Reis, M., Wernisch, L., Savva, R.: Unexpected correlations between gene expression and codon usage bias from microarray data for the whole Escherichia coli K-12 genome. Nucleic acids research 31(23), 6976–6985 (2003)

11. Duret, L.: tRNA gene number and codon usage in the C. elegans genome are co-adapted for optimal translation of highly expressed genes. Trends in Genetics 16(7), 287–289 (2000)

12. Freilich, S., Kreimer, A., Borenstein, E., Yosef, N., Sharan, R., Gophna, U., Ruppin, E.: Metabolic-network-driven analysis of bacterial ecological strategies. Genome biology 10(6), R61 (2009)

13. Gouy, M., Gautier, C.: Codon usage in bacteria: correlation with gene expressivity. Nucleic acids research 10(22), 7055–7074 (1982)

14. Grantham, R., Gautier, C., Gouy, M., Mercier, R., Pave, A.: Codon catalog usage and the genome hypothesis. Nucleic acids research 8(1), 197–197 (1980)

15. Hart, A., Cortés, M.P., Latorre, M., Martinez, S.: Codon usage bias reveals genomic adaptations to environmental conditions in an acidophilic consortium. PloS one 13(5) (2018)

16. Hershberg, R., Petrov, D.A.: General rules for optimal codon choice. PLoS genetics 5(7) (2009)

17. Hooper, S.D., Berg, O.G.: Gradients in nucleotide and codon usage along escherichia coli genes. Nucleic acids research 28(18), 3517–3523 (2000)

18. Ikemura, T.: Correlation between the abundance of Escherichia coli transfer RNAs and the occurrence of the respective codons in its protein genes: a proposal for a synonymous codon choice that is optimal for the E. coli translational system. Journal of molecular biology 151(3), 389–409 (1981)

19. Ikemura, T.: Codon usage and tRNA content in unicellular and multicellular organisms. Molecular biology and evolution 2(1), 13–34 (1985)

20. Jiang, H., Guan, W., Pinney, D., Wang, W., Gu, Z.: Relaxation of yeast mitochondrial functions after whole-genome duplication. Genome research 18(9), 1466–1471 (2008)

21. Jolliffe, I.: Principal Component Analysis. Springer Series in Statistics. Springer-Verlag (2002). DOI 10.1007/b98835

22. Kanaya, S., Yamada, Y., Kudo, Y., Ikemura, T.: Studies of codon usage and tRNA genes of 18 unicellular organisms and quantification of Bacillus subtilis tRNAs: gene expression level and species-specific diversity of codon usage based on multivariate analysis. Gene 238(1), 143–155 (1999)

23. Kudla, G., Murray, A.W., Tollervey, D., Plotkin, J.B.: Coding-sequence determinants of gene expression in Escherichia coli. Science 324(5924), 255–258 (2009)

24. Lowe, T.M., Eddy, S.R.: trnascan-se: a program for improved detection of transfer rna genes in genomic sequence. Nucleic acids research 25(5), 955–964 (1997)

25. Man, O., Pilpel, Y.: Differential translation efficiency of orthologous genes is involved in phenotypic divergence of yeast species. Nature genetics 39(3), 415–421 (2007)

26. Ollivier, B., Caumette, P., Garcia, J.L., Mah, R.: Anaerobic bacteria from hypersaline environments. Microbiology and Molecular Biology Reviews 58(1), 27–38 (1994)

27. Percudani, R., Pavesi, A., Ottonello, S.: Transfer RNA gene redundancy and translational selection in Saccharomyces cerevisiae. Journal of molecular biology 268(2), 322–330 (1997)

28. Plotkin, J.B., Kudla, G.: Synonymous but not the same: the causes and consequences of codon bias. Nature Reviews Genetics 12(1), 32–42 (2011)

29. Ran, W., Higgs, P.G.: Contributions of speed and accuracy to translational selection in bacteria. PloS one 7(12) (2012)

30. Reis, M.d., Savva, R., Wernisch, L.: Solving the riddle of codon usage preferences: a test for translational selection. Nucleic acids research 32(17), 5036–5044 (2004)

31. Rocha, E.P.: Codon usage bias from tRNA’s point of view: redundancy, specialization, and efficient decoding for translation optimization. Genome research 14(11), 2279–2286 (2004)

32. Roller, M., Lucić, V., Nagy, I., Perica, T., Vlahoviček, K.: Environmental shaping of codon usage and functional adaptation across microbial communities. Nucleic acids research 41(19), 8842–8852 (2013)

33. Salim, H.M., Cavalcanti, A.R.: Factors influencing codon usage bias in genomes. Journal of the Brazilian Chemical Society 19(2), 257–262 (2008)

34. Sharp, P.M., Li, W.H.: The codon adaptation index-a measure of directional synonymous codon usage bias, and its potential applications. Nucleic acids research 15(3), 1281–1295 (1987)

35. Sharp, P.M., Tuohy, T.M., Mosurski, K.R.: Codon usage in yeast: cluster analysis clearly differentiates highly and lowly expressed genes. Nucleic acids research 14(13), 5125–5143 (1986)

36. Sørensen, M.A., Kurland, C., Pedersen, S.: Codon usage determines translation rate in Escherichia coli. Journal of molecular biology 207(2), 365–377 (1989)

37. Tuller, T., Girshovich, Y., Sella, Y., Kreimer, A., Freilich, S., Kupiec, M., Gophna, U., Ruppin, E.: Association between translation efficiency and horizontal gene transfer within microbial communities. Nucleic acids research 39(11), 4743–4755 (2011)

38. Tuller, T., Kupiec, M., Ruppin, E.: Determinants of protein abundance and translation efficiency in S. cerevisiae. PLoS computational biology 3(12) (2007)

39. Varenne, S., Buc, J., Lloubes, R., Lazdunski, C.: Translation is a non-uniform process: effect of tRNA availability on the rate of elongation of nascent polypeptide chains. Journal of molecular biology 180(3), 549–576 (1984)

40. Vieira-Silva, S., Rocha, E.P.: The systemic imprint of growth and its uses in ecological (meta) genomics. PLoS genetics 6(1) (2010)

41. Weinstein, J.N., Myers, T.G., O’Connor, P.M., Friend, S.H., Fornace, A.J., Kohn, K.W., Fojo, T., Bates, S.E., Rubinstein, L.V., Anderson, N.L., et al.: An information-intensive approach to the molecular pharmacology of cancer. Science 275(5298), 343–349 (1997)

42. Zheng, H., Wu, H.: Gene-centric association analysis for the correlation between the guanine-cytosine content levels and temperature range conditions of prokaryotic species. BMC bioinformatics 11(11), S7 (2010)

